# Mitochondrial DNA Mutations Linking Leigh Syndrome and Carotid Atherosclerosis: A Study of Shared Genetic Pathways

**DOI:** 10.1101/2024.10.13.618117

**Authors:** Abdd Sharma, Dhruvam Shukla, Anand Kalambur Ravi, Neha Simon

## Abstract

Mitochondrial DNA encodes the genetic information necessary for mitochondrial function. In humans, mitochondrial DNA spans 16,569 base pairs while representing a small fraction of the genetic material in humans. Due to their higher mutation rates compared to nuclear DNA, Mitochondrial DNA mutations are emerging as promising biomarkers for assessing disease predisposition and progression. This in-silico analyzes the correlation between Mitochondrial DNA mutations and two conditions: Leigh syndrome, a severe neurodegenerative disorder, and carotid atherosclerosis, a major cardiovascular disease. We focus on five single nucleotide variants (SNVs) – m.14459G>A, m.13513G>A, m.12315G>A, m.1555A>G, and m.15059G>A – to explore their roles in both diseases. Our findings suggest that mutations m.14459G>A, m.1555A>G, and m.15059G>A contribute to the pathogenesis of both conditions. In contrast, m.12315G>A is linked to MELAS syndrome and carotid atherosclerosis. Interestingly, the m.13513G>A mutation is associated with Leigh syndrome but negatively correlated with atherosclerosis, suggesting a potential protective effect. These SNVs could serve as targets for diagnostic and therapeutic approaches, enhancing our understanding of the genetic basis of these diseases.

## 1. Introduction

In recent years, molecular genetic diagnostics have gained significant attention, particularly in studying polygenic multifactorial diseases such as cardiovascular pathologies and atherosclerosis[2]. Atherosclerosis, a leading cause of cardiovascular disease, primarily affects middle-aged individuals but is increasingly prevalent in younger populations[1][3][5]. This trend highlights the need for early detection methods, as conventional clinical approaches often miss atherosclerosis in its early stages[1][2]. As a result, identifying molecular genetic markers as early predictors of atherosclerosis has become a crucial research focus[2][3][4].

One promising approach is studying mitochondrial DNA (mtDNA) mutations. Mitochondrial DNA is a small circular chromosome found in the mitochondria, spans 16,569 base pairs containing 37 genes that encode for 13 proteins, 22 tRNAs, and 2 rRNAs[6][7][14][59]. Mutations that alter these base pairs, known as Single Nucleotide Variants (SNVs), occur more frequently in mtDNA than in nuclear DNA and are considered potential biomarkers for atherogenesis[6][7][14]. mtDNA may offer advantages over nuclear DNA as a biomarker for disease detection[6][14]. Each human cell contains multiple mitochondria with several copies of the mitochondrial genome, and heteroplasmy occurs when two or more mtDNA variants are present in the same cell. The level of heteroplasmy can influence the occurrence and severity of somatic mtDNA mutations, making these mutations crucial in genetic diagnostics[8][14].

Mitochondrial diseases characterized by defects in oxidative phosphorylation often result from mutations in mitochondrial genes[6][7][14]. Leigh syndrome is a severe mitochondrial neurodegenerative disorder with various neurological symptoms and mitochondrial dysfunction[9][10][11][14](Fig.1). It typically affects the MT-ATP6 gene, disrupting the ATP synthase complex, which is essential for cellular energy production[9][10][11][14]. This disease significantly reduces life expectancy to just 2 to 3 years[9][10][11][12][14]. Although rare, Leigh syndrome occurs more frequently in certain populations, such as those in Saguenay Lac-Saint-Jean, Quebec, and the Faroe Islands[9][10][11][12][14]. About 32% of Leigh syndrome cases are linked to mtDNA SNVs, underscoring the importance of molecular testing for diagnosis[9][10][11][12][13][14].

**Figure 1:**
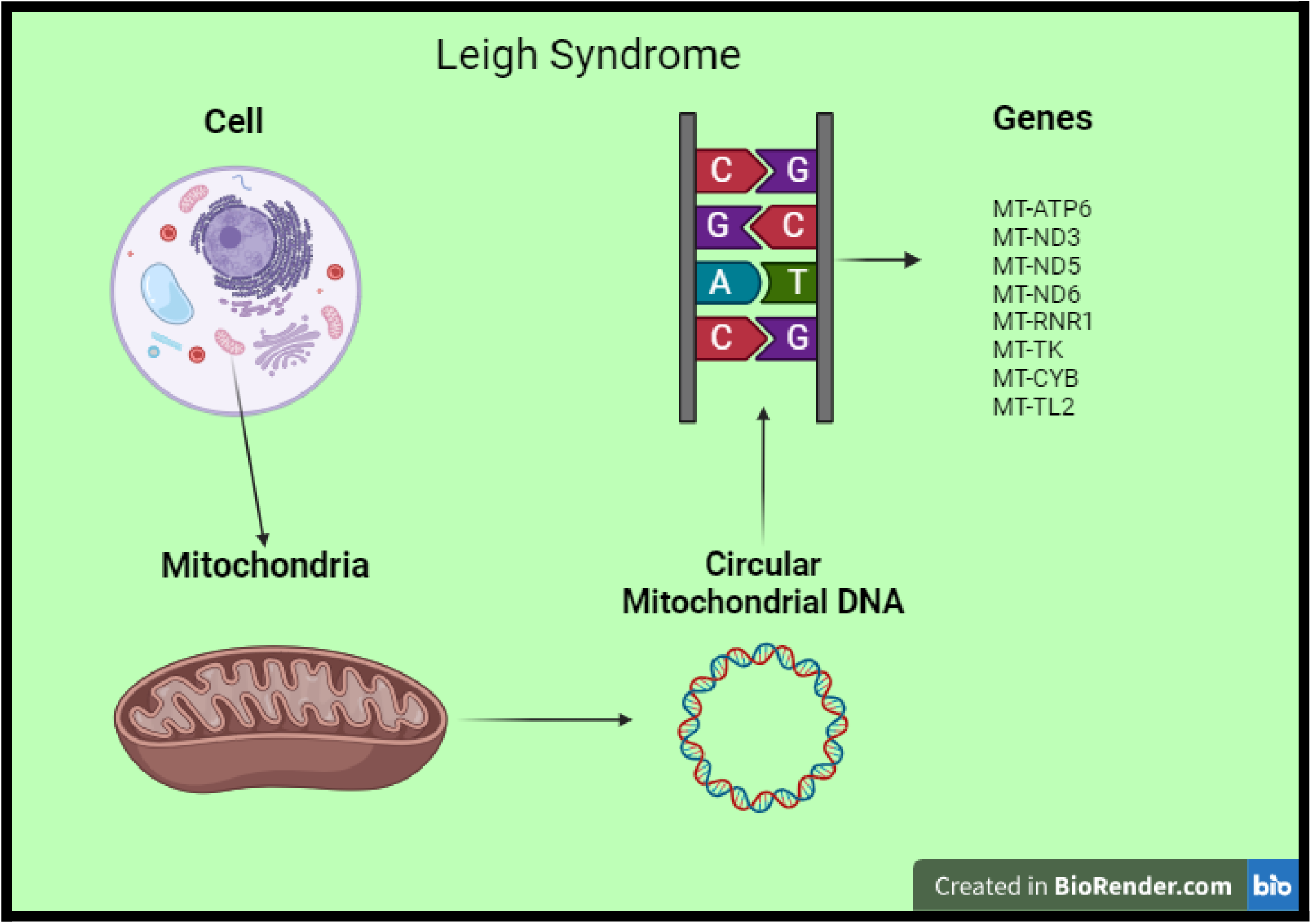
Image showcasing how Leigh Syndrome occurs due to Mitochondrial DNA mutations in certain genes from the circular DNA( Created in BioRender.com )

Carotid atherosclerosis, or carotid artery stenosis (Fig.2), is a narrowing of the blood vessels in the neck, often caused by cholesterol buildup[1][15][16][17]. This condition can lead to symptoms such as numbness, weakness, dizziness, and stroke-like events[1][15][16][17]. Without surgical intervention, carotid artery blockage significantly increases the risk of stroke, with a 23% likelihood of stroke within 5 years[1][15][16][17]. Genetic markers, particularly mtDNA SNVs, play a critical role in determining the link between mitochondrial heteroplasmy mutations and atherosclerosis[8].

**Figure 2:**
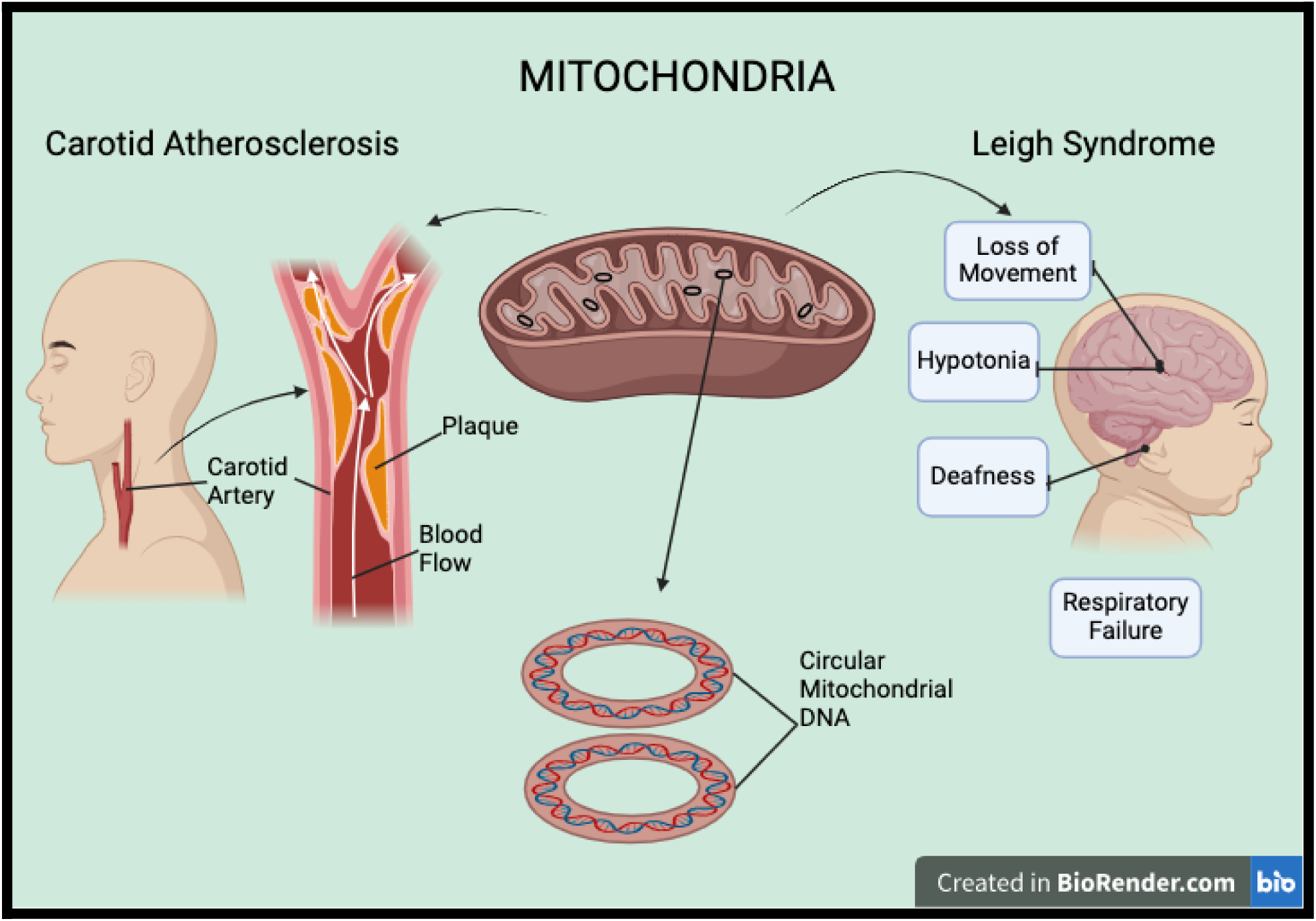
Depiction of mitochondrial circular DNA, Leigh Syndrome, and carotid atherosclerosis. Created in BioRender.com

Several SNVs have been associated with both carotid atherosclerosis and Leigh syndrome, including m.14459G>A[3][4][18][19][20][21][22][50][54], m.13513G>A[3][4][20][23][36][40], m.12315G>A[3][4][20][24][54], m.1555A>G[3][4][20][25][54], and m.15059G>A[3][4][20][26][54]. Notably, mutations such as m.14459G>A[3][4][18][19][20][21][22][50][54], m.1555A>G[3][4][20][25][54], and m.15059G>A[3][4][20][26][54] show strong associations with both conditions, suggesting these SNVs play a key role in the pathogenesis of both diseases. The m.12315G>A[3][4][20][24][54] mutation, linked to MELAS syndrome, is also associated with increased carotid atherosclerosis. MELAS syndrome, characterized by mitochondrial encephalopathy, lactic acidosis, and stroke-like episodes, shares a close relationship with Leigh syndrome[27][28]. Interestingly, the m.13513G>A[3][4][20][23][36][40] mutation, while associated with Leigh syndrome, appears to have a protective effect against atherosclerotic lesions[3][23][36][40]. This paper explores the link between mtDNA mutations and two conditions: carotid atherosclerosis and Leigh syndrome.

## 2. Materials and Methods

We systematically reviewed existing literature on Mitochondrial DNA mutations associated with Leigh syndrome and carotid atherosclerosis. Databases such as MitoVisualize, MITOMAP, GeneMANIA, and gnomAD were used to visualize and analyze genomic locations and pathogenicity of Mitochondrial DNA variants. A specific focus was placed on SNVs that are associated with both diseases. Tools like GeneMANIA were used to predict gene interactions, while MitoVisualize and MITOMAP provided insights into the disease associations of these mutations. The comprehensive flow chart of the databases and tools used to analyze the common mutation between carotid atherosclerosis and Leigh syndrome is shown in Figure 3.

**Figure 3:**
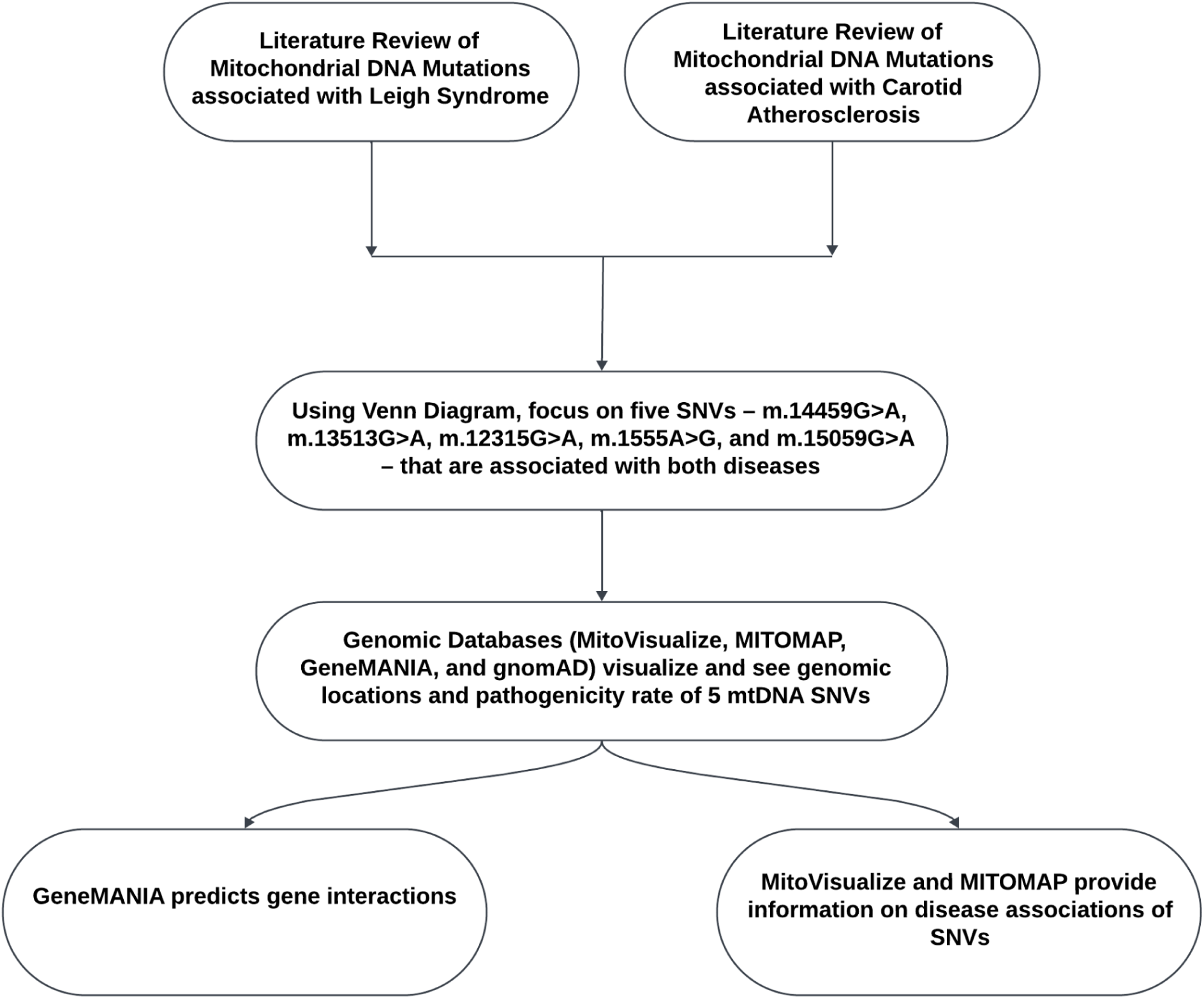
Flowchart diagram to show the material and method used to analyze the data.

## 3. Results and Discussion

First, we identified 15 mtDNA SNVs associated with Leigh syndrome, and 18 mtDNA SNVs associated with carotid atherosclerosis (*see Table 1, Table 2, and Figure 4*). A specific focus was placed on five SNVs – m.14459G>A, m.13513G>A, m.12315G>A, m.1555A>G, and m.15059G>A – that are associated with both diseases (*see Table 3 and Figure 4*).

**Figure 4:**
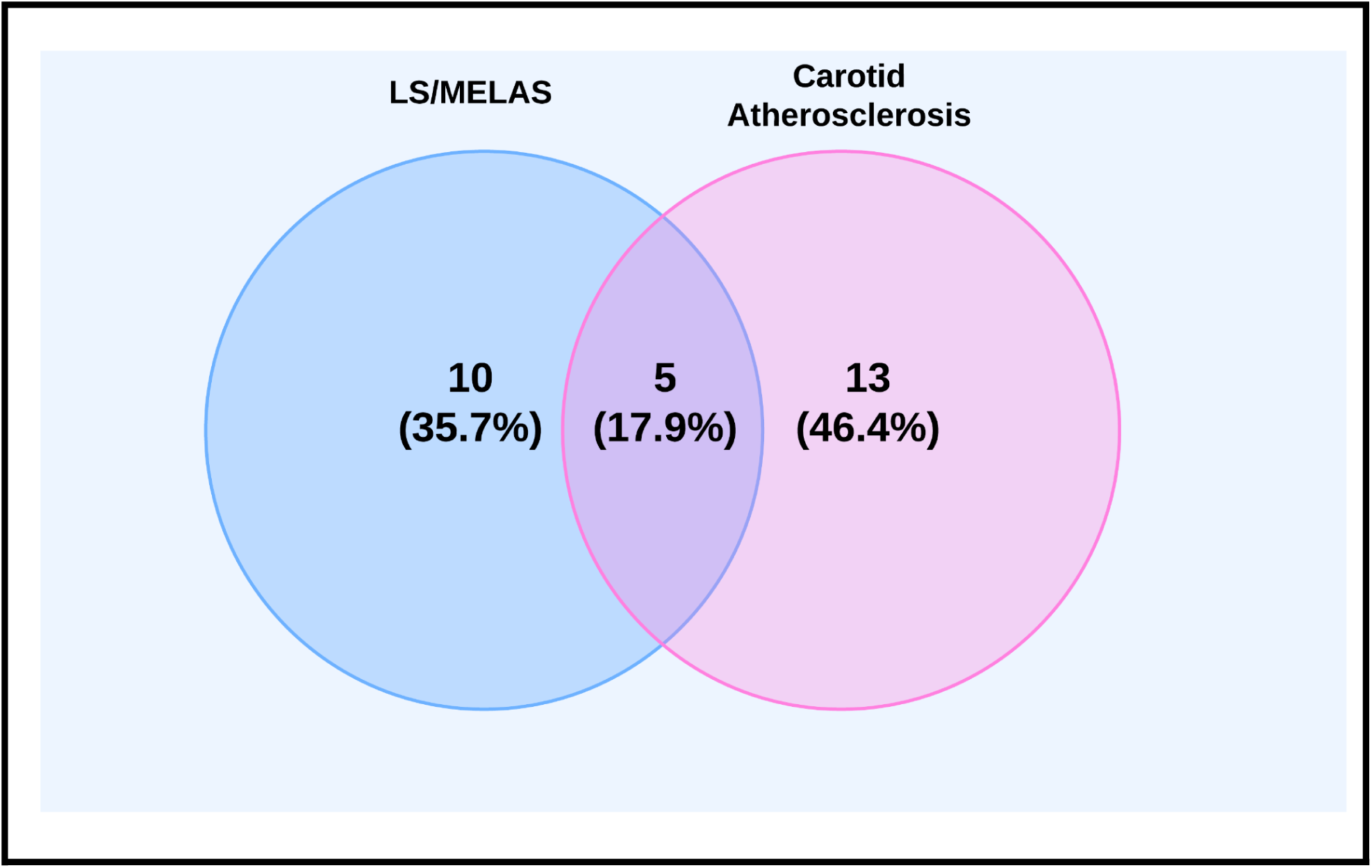
Venn Diagram of Shared and Distinct Mitochondrial DNA SNVs.The five shared SNVs are: m.14459G>A, m.13513G>A, m.12315G>A, m.1555A>G, m.15059G>A

When analyzed these SNVs using MitoVisualize, only m.12315G>A and m.1555A>G showed results[31]. This is likely because the GnomAD database requires a certain level of heteroplasmy for SNVs to be included, and only these two meet that threshold[30][31].

The m.12315G>A variant, located in the MT-TL2 gene, causes a G>A mutation in the T-stem domain of the tRNA, disrupting a Watson-Crick base pairing, indicated by the red dot in Figure 5[30][31]. This mutation is predicted to be pathogenic and is confirmed to be disease-associated[30][31].

**Figure 5:**
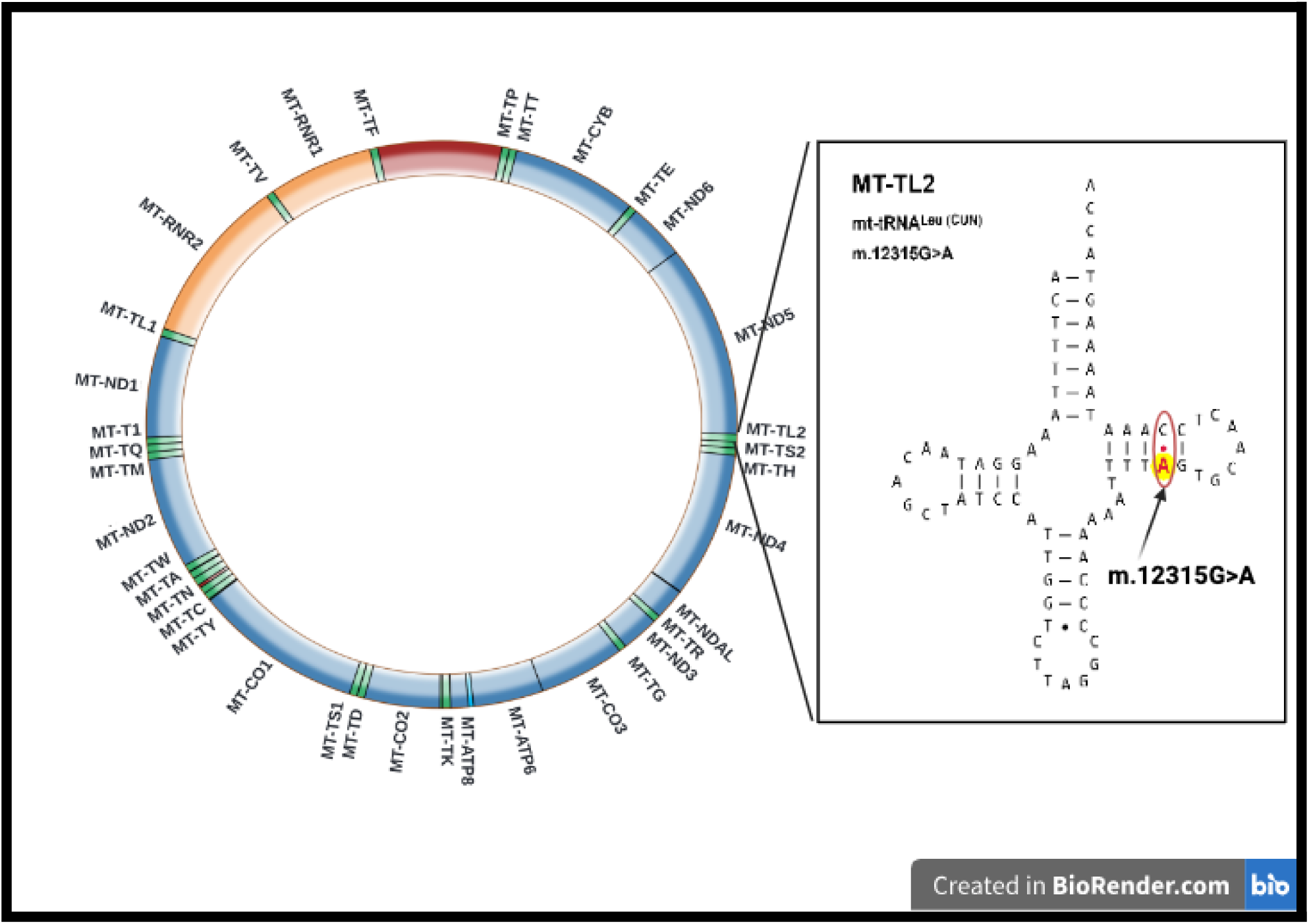
Visualization of m.12315G>A SNV, showing that it is found in MT-TL2 gene in the mitochondria and that it codes for tRNA, disrupting a Watson-Crick base pairing. Created in BioRender.com

**Figure 6:**
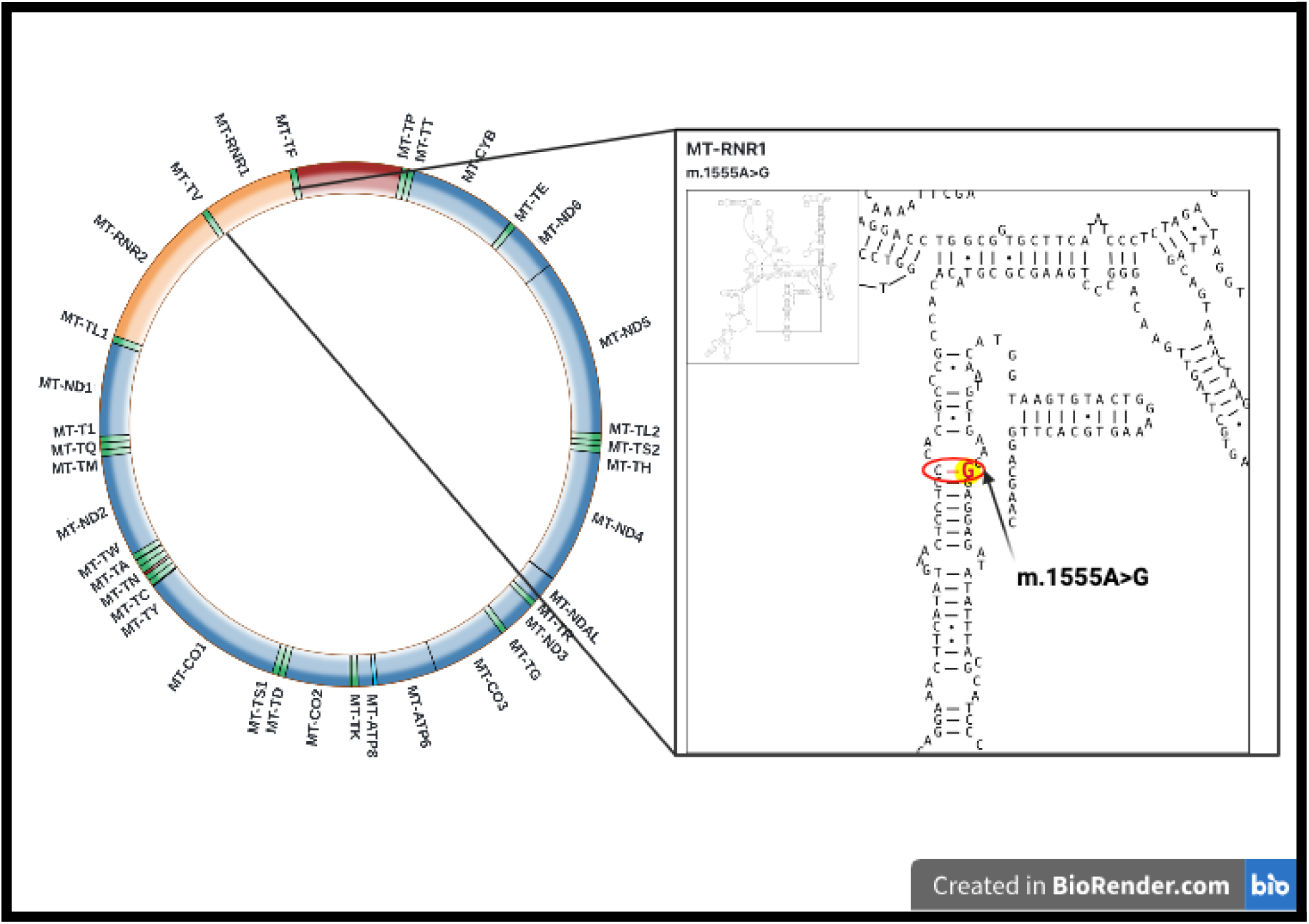
Visualization of m.1555A>G SNV, showing that it is found in MT-RNR1 gene in the mitochondria and that it codes for rRNA, adding a Watson-Crick base pairing. Created in BioRender.com

The m.1555A>G variant in the MT-RNR1 gene leads to an A>G mutation in the rRNA, forming a Watson-Crick base pairing, represented by the red line in Figure 6[30][31]. This mutation is also confirmed to be associated with disease[30][31].

We further visualized all five SNVs in the circular mitochondrial genome using MitoVisualize (Fig. 7)[30][31].

**Figure 7:**
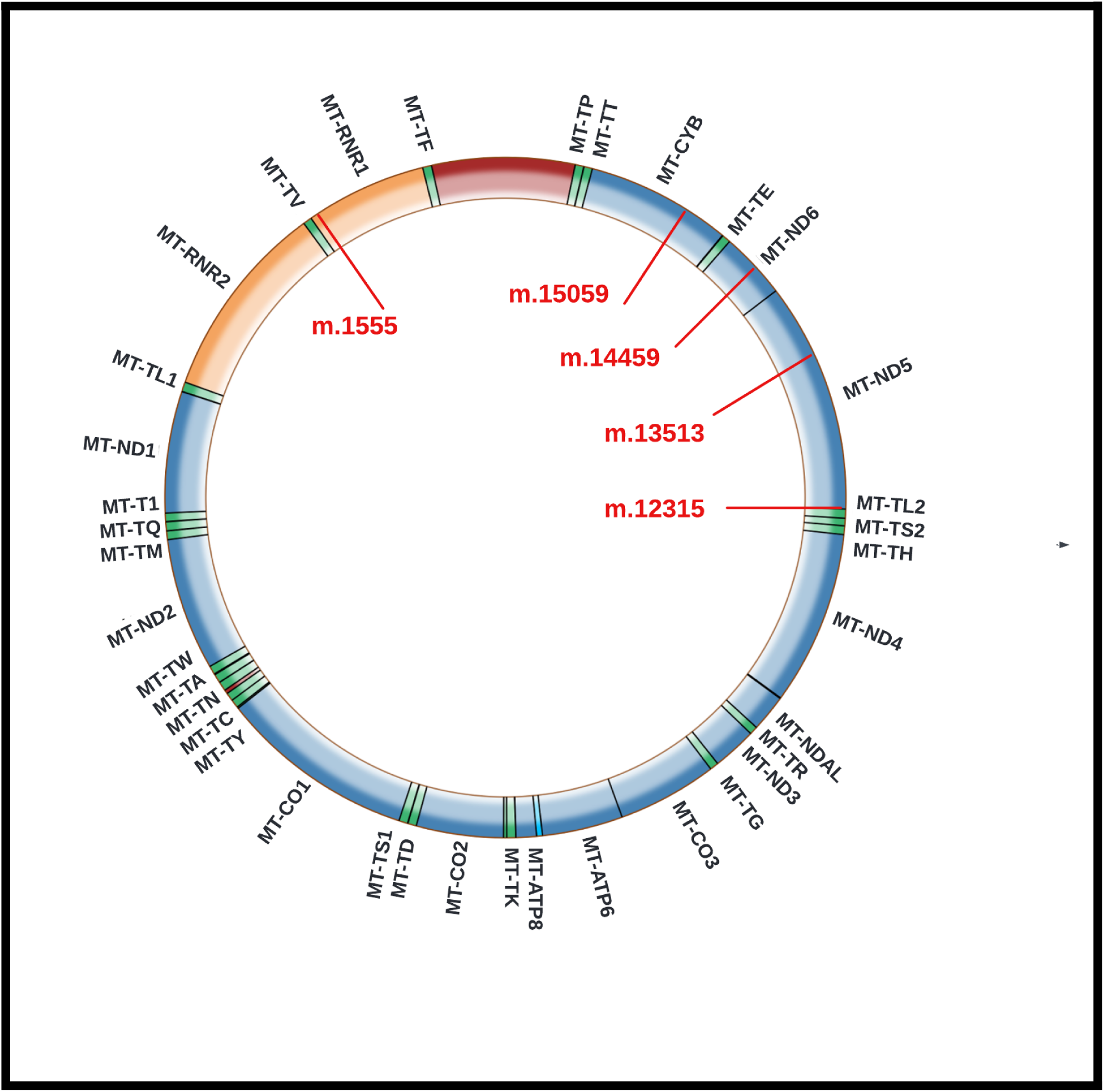
All 5 SNVs displayed in the circular mitochondrial genome

Additionally, an analysis using MITOMAP revealed that m.14459G>A is associated with both Leigh syndrome and carotid atherosclerosis, m.13513G>A is linked to Leigh syndrome/MELAS and negatively associated with carotid atherosclerosis, m.12315G>A is associated with both MELAS and carotid atherosclerosis, m.1555A>G is possibly atherosclerotic, and m.15059G>A is linked to both Leigh syndrome and carotid atherosclerosis[33].

The SNVs were located within specific mitochondrial genes: m.12315G>A in MT-ND4, m.13513G>A in MT-ND5, m.14459G>A in MT-ND6, and m.15059G>A in MT-CYB[33].

Using a GeneMANIA graph (Fig. 8), we identified several genes interacting with MT-ND1, MT-ND4, MT-ND5, MT-ND6, and MT-CYB, including MT-ND2, MT-CO1, MT-CO2, NDUFA8, and others[34]. About 77.64% of these networks are based on physical interactions, suggesting that pathogenic mutations in these SNVs could affect these genes[34]. Another 8.01% of the networks involve co-expression, which may link these genes to the expression of Leigh syndrome or carotid atherosclerosis[34].

**Figure 8:**
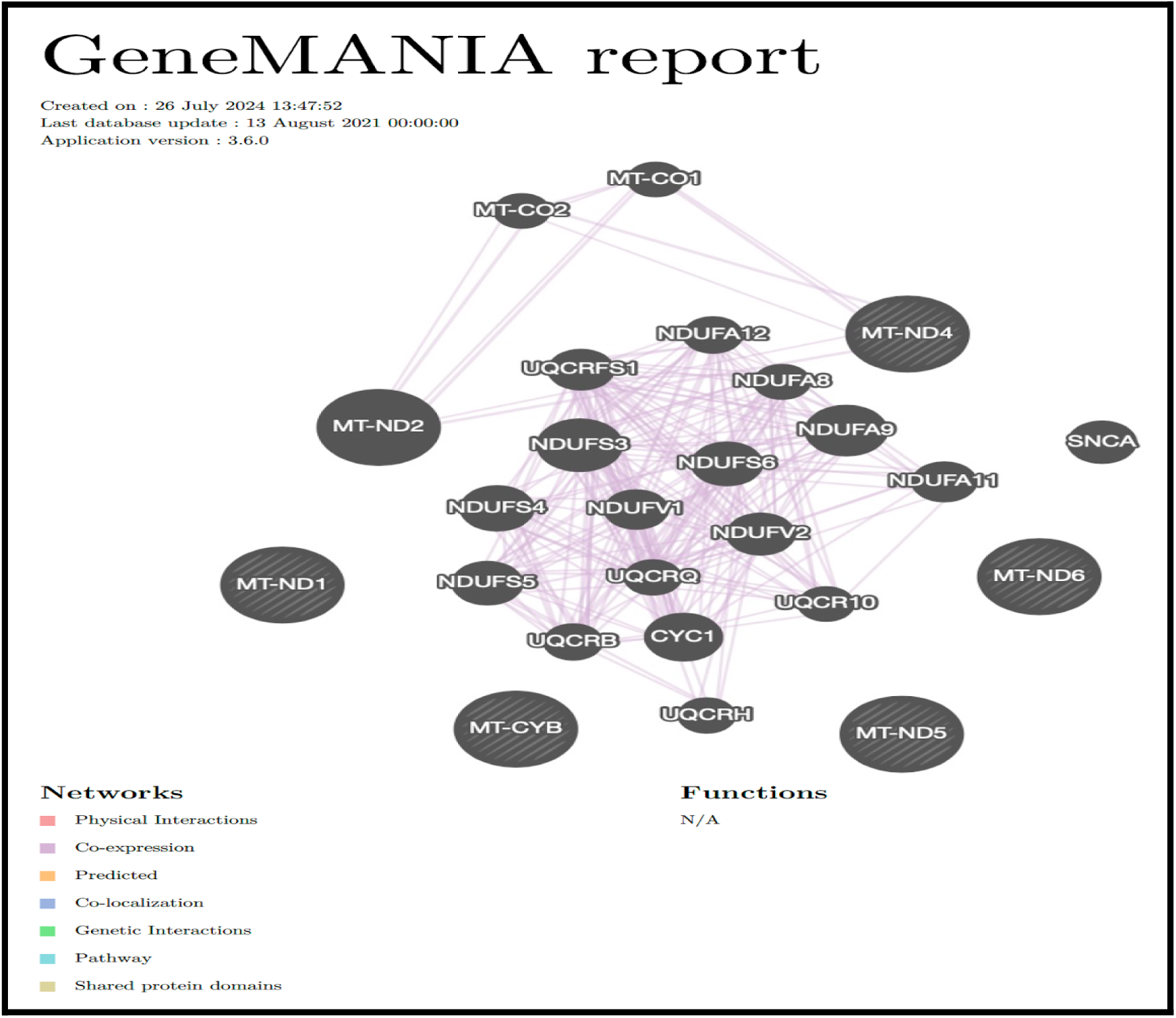
GeneMANIA graph with the coexpression (in purple) between genes in which the SNVs are present[34].

Finally, we present a case study of a 47-year-old female patient with carotid artery stenosis, primarily due to carotid atherosclerosis, and Leigh-like syndrome. This case underscores the clinical relevance of these findings and highlights the shared genetic underpinnings of these conditions, providing insights into their pathophysiology and opening avenues for novel therapeutic strategies[35]

**Table 1:** 15 Mitochondrial DNA Single Nucleotide Variants associated with Leigh Syndrome.

| **mtDNA Mutation Associated with Leigh Syndrome** | **Location in Mitochondrial Circular Genome** | **Pathogenicity** | **References** |
| --- | --- | --- | --- |
| m.9185T>C | MT-ATP6 | Likely Pathogenic | [3], [4], [20], [23], [36], [40] |
| m.9176T>C | MT-ATP6 | Pathogenic | [42], [47], [51], [52] |
| m.8993T>G | MT-ATP6 | Like Pathogenic | [40], [42], [47], [49], [51], [52] |
| m.8344A>G | MT-TK | Pathogenic | [42], [51] |
| m.15059G>A | MT-CYB | Likely Pathogenic | [3], [4], [20], [26], [54] |
| m.13513G>A | MT-ND5 | Pathogenic | [40], [42], [47], [51], [52] |
| m.13046T>C | MT-ND5 | Likely Pathogenic | [40] |
| m.9176T>G | MT-ATP6 | Pathogenic | [42], [47], [51] |
| m.3243A>G | MT-RNR1 | Pathogenic | [37], [42], [47], [51], [53] |
| m.14459G>A | MT-ND6 | Likely Pathogenic | [3], [4], [18], [19], [20], [21], [22], [50], [54] |
| m.8993T>C | MT-ATP6 | Pathogenic | [40], [42], [51], [47], [52] |
| m.10191T>C | MT-ND3 | Pathogenic | [42], [47] |
| m.8597T>C | MT-ATP6 | Pathogenic | [52] |
| m.1555A>G | MT-RNR1 | Pathogenic | [3], [4], [20], [25], [54] |
| m.12315G>A | MT-TL2 | Pathogenic | [3], [4], [20], [24], [54] |

**Table 2:** 18 mitochondrial DNA single nucleotide variants associated with Carotid Atherosclerosis.

| mtDNA Mutation Associated with Carotid Atherosclerosis | Location in Mitochondrial Circular Genome | Pathogenicity | References |
| --- | --- | --- | --- |
| m.652delG | MT-RNR1 | Likely Pathogenic | [3], [4] |
| m.3336T>C | MT-ND1 | Reported Pathogenic | [3], [4] |
| m.14459G>A | MT-ND6 | Likely Pathogenic | [3], [4], [18], [19], [20], [21], [22], [50], [54] |
| m.15059G>A | MT-CYB | Likely Pathogenic | [3], [4], [20], [26], [54] |
| m.14798T>C | MT-CYB | Likely Pathogenic | [4] |
| m.3336C>T | MT-ND1 | Likely Pathogenic | [3] |
| m.1555A>G | MT-RNR1 | Pathogenic | [3], [4], [20], [25], [54] |
| m.5178C>A | MT-ND2 | Reported Pathogenic | [3], [4], [54] |
| m.13513G>A | MT-ND5 | Pathogenic | [3], [4], [20], [23], [36], [40] |
| m.576insC | MT-CR (control region) | Likely Pathogenic | [4], [54] |
| m.14798C>T | MT-CYB | Likely Pathogenic | [3], [4], [54] |
| m.12372G>A | MT-ND5 | Reported Pathogenic | [3], [4], [20], [24], [54] |
| m.3256C>T | MT-TL1 | Pathogenic | [3], [4] |
| m.12315G>A | MT-TL2 | Pathogenic | [54] |
| m.14846G>A | MT-CYB | Reported Pathogenic | [4], [54] |
| m.3480A>G | MT-ND1 | Likely Pathogenic | [54] |
| m.13708G>A | MT-ND5 | Pathogenic | [3], [4], [54] |
| m.15326A>G | MT-CYB | Likely Pathogenic | [3], [4], [20], [24], [54] |

**Table 3:** 5 shared mitochondrial DNA single nucleotide variants associated with Carotid Atherosclerosis and Leigh Syndrome.

| mtDNA Mutation Associated with both diseases | Location in Mitochondrial Circular Genome | Pathogenicity | References |
| --- | --- | --- | --- |
| m.14459G>A | MT-ND6 | Likely Pathogenic | [3], [4], [18], [19], [20], [21], [22], [50], [54] |
| m.15059G>A | MT-CYB | Likely Pathogenic | [3], [4], [20], [26], [54] |
| m.1555A>G | MT-RNR1 | Pathogenic | [3], [4], [20], [25], [54] |
| m.13513G>A | MT-ND5 | Pathogenic | [3], [4], [20], [23], [36], [40] |
| m.12315G>A | MT-TL2 | Pathogenic | [54] |

## 4. Conclusion

This study has identified five Mitochondrial DNA SNVs – m.14459G>A, m.13513G>A, m.12315G>A, m.1555A>G, and m.15059G>A – that are linked to both Leigh syndrome and carotid atherosclerosis. Mutations m.14459G>A, m.1555A>G, and m.15059G>A are particularly significant, as they are associated with both conditions. In contrast, m.13513G>A may have a protective effect against atherosclerosis. These findings suggest that Mitochondrial DNA mutations may serve as biomarkers for disease progression and provide new avenues for the development of gene therapies targeting mitochondrial dysfunctions common to both diseases.

Future research should focus on exploring the heteroplasmy and tissue distribution of these mutations and developing therapeutic strategies that address the shared mitochondrial dysfunctions. By understanding the complex interplay between genetic mutations and disease, we can advance our knowledge of the molecular mechanisms underlying Leigh syndrome and carotid atherosclerosis, paving the way for innovative treatment options.

